# Finite-State Likelihood: an Approximation for Genome Phylogeny Inference based on Generalized Gene Contents

**DOI:** 10.64898/2025.12.08.693040

**Authors:** Xun Gu

## Abstract

Rapid growth of entire genome data has revolutionized the field of phylogenomics, i.e., the problem of tree of life. Substantial studies demonstrated that genome phylogeny can be inferred based upon the generalized gene content approach. Two simple types were widely-used: the first-order gene content (J=1) for the presence or absence of a gene family, and the second-order gene content (J=2) for the extended gene content (absence, single-copy, or duplicates). Moreover, a specific form of birth-death-input process was invoked to model the evolutionary process of a gene family, taking gene duplication, gene loss and new gene origin or horizontal gene transfer into account.

*Gu X. Genome distance and phylogenetic inference accommodating gene duplication, loss and new gene input, Mol Phylogenet Evol 2023]*.

Though genome distance methods have been successful for genome phylogeny inference, the maximum likelihood (ML) approach is subject to a huge computation burden. In this article, I formulate a finite-state ML approximation to solve this problem. For a given J-order gene contents, the evolution of a gene family along a phylogeny is modeled by a stochastic process with a finite (J+1) number of states. Consequently, the computational cost of a finite-state likelihood for a given phylogeny is comparable to a typical sequence-based likelihood function. Two analyses were carried out as a proof of concept, including a simulation study to examine the performance of phylogenetic inference, and a case study to evaluate to what extent the Fixed-State ML can be used to determine the root of the genome phylogeny. Overall, the Fixed-State ML may shed lights on the feasibility of phylogenetic likelihood analysis on the pattern of genome evolution.

## Introduction

Phylogenomics focuses on analyzing evolutionary histories and reconstructing the relationships between taxa in a broad range of taxonomic hierarchy [1-6]. For decades, researchers have approached to the inference problem of genome phylogeny based upon the size variation of gene families, or the generalized gene concepts. Under the phylogeny of organisms, the genome process can be intuitively viewed as follows [7]: a gene family in an organism was generated by a single event of new gene input (new gene origin or horizontal gene transfer), expanded by one or more rounds of gene duplications, and perhaps reduced or even lost as the result of null mutations (being pseudogenes). There are many reported approaches, which can be unified under the concept of generalized gene content. For instance, the first-order gene content (J=1) refers to the presence or absence of a gene family in a genome [8-12]. Meanwhile, the second-order gene content (J=2) refers to the extended gene content (absence, presence as a single-copy, and duplicates) [13, 14]. Those methods, explicitly or implicitly, invoked the birth-and-death process to model the evolutionary process of gene families, which recently was further extended to the inclusion of the mechanism of new gene input [14].

Many statistical methods have been implemented for practical analysis [10, 11, 13-20]. It should be noticed that most of them aimed at the estimation of genome distance between two species. The performance of genome distance approach is generally satisfactory under the birth-death process with J=1 or J=2 [13]. However, [14] found that an accurate estimation of genome distance was difficult when the mechanism of new-gene input was taken into account, indicating that the pairwise distance approach may not be sufficient to account for the model complexity of genome evolution. Therefore, sophisticated methods such as maximum likelihood are desirable.

Zhang and Gu [11] formulated a maximum likelihood in the case of four genomes under a simple case (J=1). However, the algorithm of Zhang and Gu [11] encountered two technical difficulties. First, when a sophisticated model is implemented, i.e., J=2, the analytical form of the likelihood would be too tedious to be useful in practice. Second, when more-than-four genomes are considered, the computational cost would increase exponentially. The underlying reason can be attribute to the ‘infinite-state’ problem: for each likelihood function derived by Zhang and Gu [11], one has to calculate the very large sum (as an approximation for infinity) at each internal node, resulting in a huge computational burden when the phylogeny is large.

The finite-state approximation of maximum likelihood (Finite-State ML for short) aims at the reduction of computational cost. The Finite-State ML postulates that, for any J-th order gene content, a gene family at any internal node has a finite number of states about its size, which are 0 (no gene), 1 (single gene), 2 (two duplicates) until the last state (equal to J or more number of member genes). One may anticipate that the Finite-State ML has two advantages. First, the computational cost of Finite-State ML is generally comparable to that of sequence-based Markov-chain likelihood function along the same phylogeny. Many ML algorithms originally developed for DNA sequences can be applied to the Finite-State ML after some technical modifications. Second, the computation time is roughly linear to the model complexity, either a higher order of gene contents or the evolutionary model, making the hierarchical likelihood ratio test feasible to investigate the pattern of genome evolution. The challenge is, to implement Finite-State ML, one must renormalize the original transition probabilities (with an infinite number of states) to those with a finite (J+1) states. To this end, I design a practically feasible procedure of renormalization. As a proof of concept, some analyses were carried out in the case of four genomes: the simulation study to examine the performance of phylogenetic inference, and the case study to explore to what extent the Fixed-State ML can be used to determine the root of the genome phylogeny. The challenges of the Fixed-State ML on detecting the genome phylogeny and the pattern of genome evolution are discussed.

## Methods

### The birth-death-input model of gene family size evolution

Whole-genome comparison has revealed a substantial size variation of gene families across genomes, highlighting the evolutionary dynamics of a gene family that can be generated (new gene input), expanded (gene duplication), and lost (being pseudogenes). Gu and Zhang [13] developed a birth-death stochastic model that included two major evolutionary processes (gene loss and gene duplication), as introduced below briefly (Table 1).

**Table 1.** A summary of important model parameters and interpretations.

| Model parameter | Interpretation |
| --- | --- |
| $\lambda$ | Evolutionary rate of gene proliferation due to gene duplication |
| $\mu$ | Evolutionary rate of gene loss |
| $\rho$ | Evolutionary rate of new gene input |
| $\delta=\rho/\lambda$ | Coefficient of new gene input |
| $\alpha$ | Proliferation parameter due to gene duplication, defined by Eq.(2) |
| $\beta$ | Loss parameter, defined by Eq.(2) |
| $J$ | The order of gene content information |
| $J=1$ | The binary gene content information (present or not present) |
| $J=2$ | The extend gene content information (single copy, duplicate, or not present) |

Let *μ* be the evolutionary rate of gene loss and *λ* be that of gene proliferation by duplication, respectively. Suppose that a gene family has *r* member genes at *t*=0. If each gene is subject to the same chance to be lost or duplicated, it has been shown that the number of member genes after *t* time units, denoted by *X*_*t*_, satisfies the following distribution

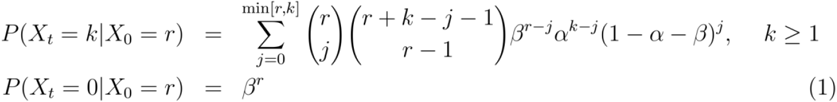

where the proliferation parameter *α* and the loss parameter *β* are given by

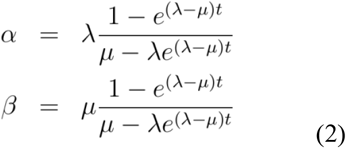

respectively, where *α*/*β*= *λ/μ* is called the proliferation-loss (*P/L*) ratio. Note that the size of gene family under the birth-death model is expected to be *X*_*0*_*e*^*-(λ-μ)t*^. Eq.(2) suggests that *α*>*β* (or *P/L*>1) indicates, on average, an increase of gene family size during the evolution, and *vice versa*.

The stochastic model of gene family size evolution was recently extended to the case that involves the new gene input (origin of new gene or lateral gene transfer). Gu [14] used a linear birth and death process, namely, the average numbers of births (*λ*_*i*_) and deaths (*μ*_*i*_) per gene family in the genome are proportional to the number of existed member genes, respectively. With the new gene input, one may write

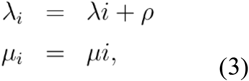

where λ and μ are the birth (gene duplication) and death (gene loss) rates, respectively, and *ρ* is the rate of new gene input [21, 22]. If *ρ*=0, the model is reduced to the pure birth and death process.

Let *δ* be the coefficient of new gene input, defined by the ratio of new gene input rate to the birth (duplication) rate, that is,

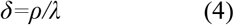

Putting together, Gu [14] showed

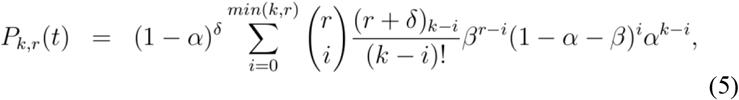

where (*r+δ*)_*k-i*_=(*r+δ*)×(*r+δ+*1)… ×(*r+δ+k-i-*1).

Eq.(5) works for any case of *λ*≠*μ*. It appears that the case of equal gene birth (duplication) and loss rates, i.e., *λ=μ*, needs some special treatments: the transition probability in this case is given by

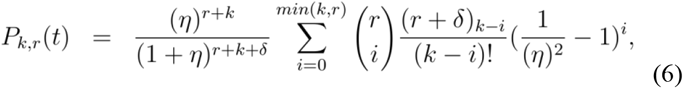

where *η*=*λt=μt*.

### The concept of generalized gene content

Based on the transition probability given by Eq.(5) that includes Eq.(1) and Eq.(6) as special cases, it is theoretically not difficult to build a likelihood function for phylogenetic analysis, as illustrated by a two-genome case. Consider two species that have been diverged *t* time units ago (Fig.1). For a given gene family that has *r* member genes at *t*=0 (in the common ancestor), there are *X*_*1*_ and *X*_*2*_ number of member genes in each genome, respectively. Under the assumption of independent evolution between lineages, the (conditional) joint probability is simply given by *P*(*X*_*1*_, *X*_*2*_|*X*_*0*_*=r*)= *P*(*X*_*1*_|*X*_*0*_*=r*)×*P*(*X*_*2*_|*X*_*0*_*=r*). Since the size of a gene family in the ancestral genome is unknown, a (prior) distribution for *X*_*0*_*=r* is assumed, denoted by *π*(*r*). Hence, the joint probability of *X*_*1*_ and *X*_*2*_ is given by

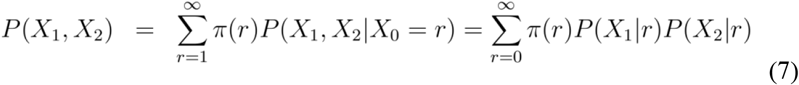

**Fig. 1.**
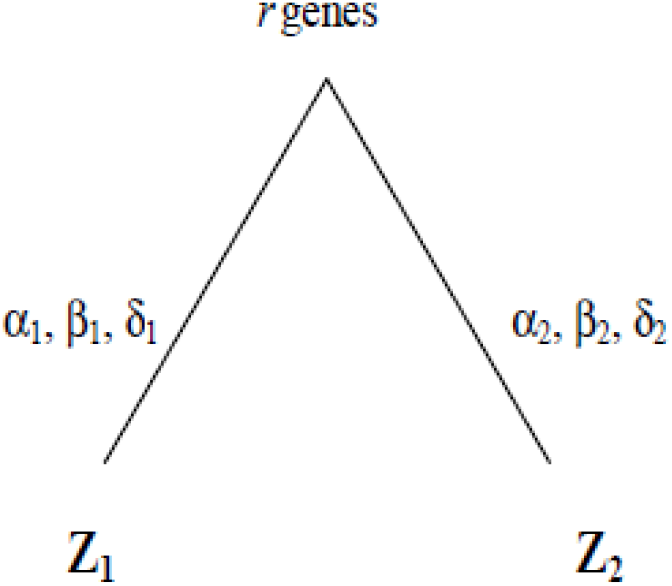
A tree of two genomes under study, where *λ*_*i*_, *μ*_*i*_, *δ*_*i*_, are the corresponding parameters of gene duplication, gene loss and new gene input in each lineage, *i*=1, 2.

Note that the complexity of transition probability in Eq.(7) makes it computationally difficult for implementation: it becomes virtually impossible even for four species [11]. Several methods were then proposed to address this issue, entitled by the concept of generalized gene content, or the J-th order gene content. The first-order gene content (J=1) refers to the simple presence-absence of a gene family in a genome, which has been widely used [8-12]. While the first-order gene content is useful for genome phylogeny inference, it seems that the binary gene-content may not have information sufficient to explore the pattern of genome evolution.

Meanwhile, the second-order gene content (J=2) refers to the extended gene content (absence, single-copy, or duplicates) [13, 14]. Here we use the birth-death-new input model as example. Let *Z*=0, 1 or 2 be the gene content index of J=2 for a gene family that has zero, one, and two-or-more member genes, respectively. According to Eq.(5), the transition probabilities of J-th order gene content can be concisely written as follows

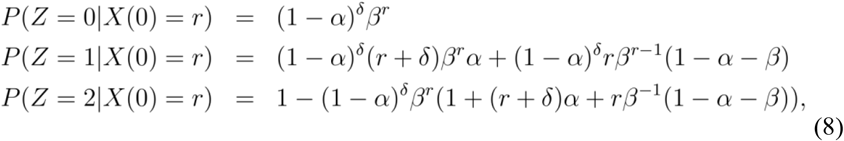

In summary, the generalized gene content can be constructed as follows. For any J-th order gene content, there are J+1 gene context indexes: for *Z*=0, 1, …, *J*-1, each of them corresponds to the number of member genes of a family, while *Z=J* corresponds to the state that a gene family has *J* or more member genes. In a broad sense, one may also group some adjacent states corresponding to a single gene content index. For instance, one may construct the third order gene content (J=3) as follows: *Z*=0 for zero gene of a family, *Z*=1 for single gene, *Z*=2 for a small gene family, and *Z*=3 for a large gene family; the criteria to distinguish between small and large gene families can be empirically specified.

### The infinite-state formulation

Given the J-th order gene content, the likelihood function under a given phylogeny can be formulated with respect to the gene content *Z*-index that has a pre-specified, fixed (J+1) number of states. However, a rigorous derivation of the likelihood requires a sum of each internal node over all (infinite) possible numbers of member genes. In the case of two genomes, for instance, let *Z*_*1*_ and *Z*_*2*_ be the gene content indexes of genome 1 and genome 2, respectively, *Z*_*1*_, *Z*_*2*_=0,…, *J*. Similar to Eq.(7), the joint probability of *Z*_*1*_ and *Z*_*2*_ is given by

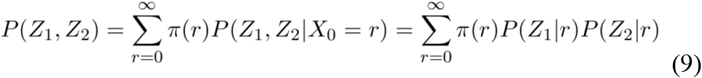

The term ‘infinite-state formulation’ is used because one has to calculate the sum at the internal node (the root), yet it is generally convergent. Given the geometric distribution *π*(*r*)=(1-*f*)^*r*^*f*, Gu and Zhang [13] obtained the analytical form of *P*(*Z*_*1*_, *Z*_*2*_) under J=2, and Zhang and Gu [11] obtained the analytical form of four genomes, *P*(*Z*_*1*_, *Z*_*2*_, *Z*_*3*_, *Z*_*4*_), under J=1. Those previous studies have shown that the calculation in general is too cumbersome to be useful except for simple cases.

### The finite-state approximation

The finite-state approximation postulates that, given the J-th order gene content, a gene family at any internal node of the phylogeny has a finite number (*J*+1) of states, which are 0 (no gene), 1 (single gene), 2 (two duplicates), until the last state (J or more member genes). To implement this approach, one must renormalize the original transition probabilities given by Eq.(5) (the birth-death process with new gene input) such that the renormalized transition probabilities can be written by a (J+1) × (J+1) matrix. The renormalization process is exemplified in the following by the second-order gene content (J=2). Let 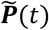 be the finite-state (3 × 3), renormalized transition probability matrix, *i, j*=0, 1 or 2, as defined by

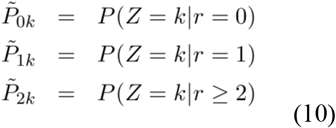

where *k*=0, 1, or 2, respectively. It appears that 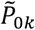 and 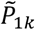 are directly from the original transition probabilities as shown by Eq.(8), whereas 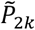 needs to be renormalized, conditional of *r* ≥ 2. Let *P*(*r* ≥ 2) be the (prior) probability of being the state of *r* ≥ 2. Under the geometric distribution *π*(*r*), one can show *P*(*r* ≥ 2) = (1 − *f*)^2^. By the basic rule of probability, we have

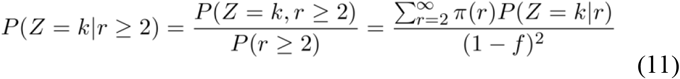

Plugging the analytical results of *P*(*Z*|*r*) in Eq.(8) onto Eq.(10) and Eq.(11), one can obtain the analytical results of the finite-state renormalized transition probabilities as follows

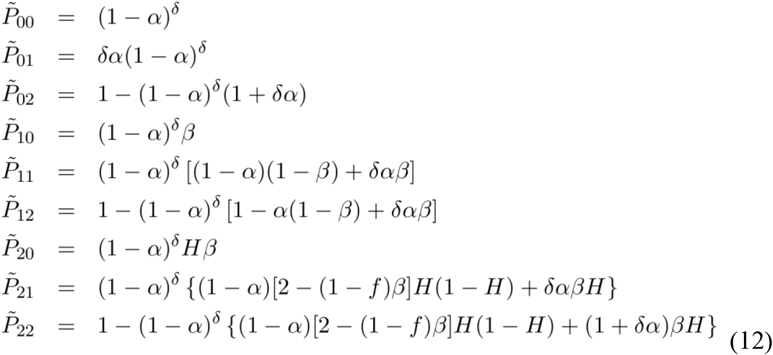

respectively, where the parameter *H* is given by

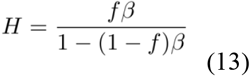

Thus, by the finite-state approximation, the joint probability of *Z*_*1*_ and *Z*_*2*_ can be written as a sum of three components, that is,

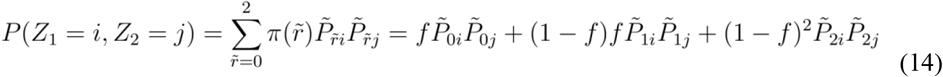

where *i, j*=0, 1, or 2.

## Results

### The finite-state renormalized transition probability matrix

Following the procedure described above, one can formulate a general likelihood framework for a given *J*-th order gene content, which have *J*+1 states. Let 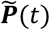 be the finite-state (*J* + 1) × (*J* + 1) renormalized transition probability matrix, whose elements are represented by 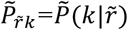. The finite-state approximation postulates that the transition probability 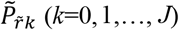 is given by

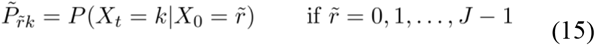

where the right hand of Eq.(15) is given by Eq.(5) (the birth-death process with new gene input), and if 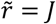 it is given by

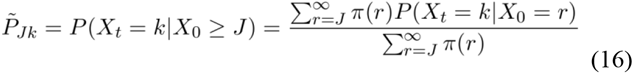

If *π*(*r*) follows a geometric distribution, one can show that 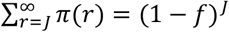.

### Likelihood under the finite-state approximation

Following the procedure described above, one can formulate a general likelihood framework for a given *J*-order gene contents, which have *J*+1 states. Here I use four genomes as an example to derive the finite-state joint distribution for the J-th order of gene contents. There are 15 possible rooted phylogenetic trees, which can be further divided into two different phylogenetic structures as shown by Fig.2. First consider the tree structure of Fig.2(a). For a given gene family, suppose there are *r*_*0*_ member genes at *t*=0 (the common ancestor), as well as *r*_*A*_ and *r*_*B*_ member genes at the two internal nodes, respectively. Under the finite-state approach, we have *r*_*0*_, *r*_*A*_, *r*_*B*_=0,…, *J*, respectively. Let *Z*_*i*_, *i*=1,…, 4, be the gene content index in genome *i*. We use *α*_*k*_, *k* = 1,…, 6, to denote the proliferation parameter in each branch, as well as *β*_*k*_ to denote the loss parameter, and *δ*_*k*_ for the coefficient of new gene input, respectively. To be concise, all those parameters are represented by a parameter vector ***Φ***. Under the assumption of independent evolution between lineages, the joint probability for any J-th order gene contents in four genomes is given by

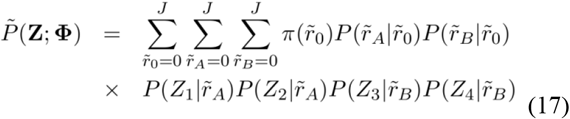

**Fig. 2.**
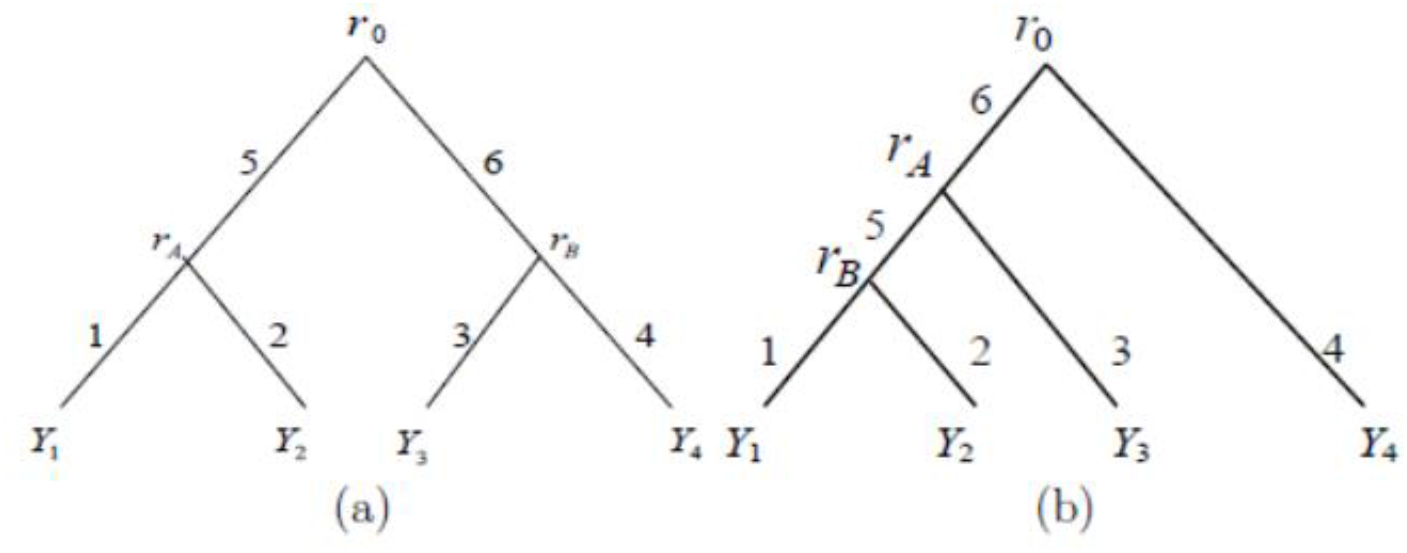
Two type of rooted phylogeny four genomes considered in the simulation study: (a) symmetric pattern and (b) asymmetric pattern. The numbers, 1, 2, …, 6, in the figure represent different branches. *Y*_*1*_, *Y*_*2*_, *Y*_*3*_ and *Y*_*4*_ are the numbers of gene families in the current genomes, respectively. Numbers of gene families at the root, and two internal nodes are denoted by *r*_*0*_, *r*_*A*_ and *r*_*B*_.

The likelihood function for Fig.2 (b) can be derived in the same way, that is,

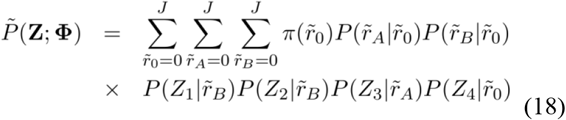

Note that the number of gene families that have zero members in all species (i.e., ***Z***=**0**) is unobservable, which should be removed from the likelihood function. Thus, the finite-state ML function for the *k*-th gene family is given by

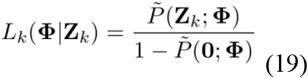

It appears that, for *m* gene families under study, the genome-wide likelihood function can be generally written as follows

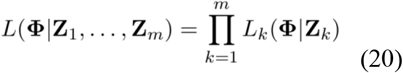

One may envisage two advantages of the Finite-State ML. First, the computational cost of the Finite-State ML (such as J=3) is generally comparable to the sequence-based likelihood along a similar phylogeny. Many ML algorithms originally developed for DNA sequences can be applied to the Finite-State ML after some technical modifications. Second, the computation time increases with the increasing of model complexity; for instance, the computational cost of J=19 is roughly equivalent to the amino acid sequence likelihood. Moreover, the maximum likelihood ratio test is generally feasible to explore the pattern of genome evolution. However, the parameter space of finite-state ML is much more complex than the sequence ML. It remains a great challenge to develop an efficient algorithm for searching the optima.

### Simulation study

I used a simulation study to address whether the finite-state ML has a similar performance as the accurate infinite-state ML. Because of the computational cost, the simplest model (*J*=1 with the pure birth-death process) in the case of four genomes was chosen as the simulation platform. Three methods, the infinite-state ML (Zhang and Gu 2004), the finite-state ML (currently) and the genome distance method (Gu and Zhang 2004), were compared.

The (rooted) phylogenetic tree was simulated based on Fig.2(a) or (b), respectively. Since the simulation results were virtually the same between them, in the following we only present the results based on the phylogeny of Fig.2(a). Three simulation themes are considered; see Table 2 for branch lengths of gene loss ***μt***=(*μt*_*1*_, *μt*_*2*_, *· · ·, μt*_*6*_), branch lengths of gene proliferation ***λt***=(*λt*_*1*_, *λt*_*2*_,*…, λt*_*6*_), as well as the corresponding parameters *α*_*i*_, *β*_*i*_ (*i*=1, 2,…, 6) calculated by Eq.(2). Branch lengths of gene proliferation and gene loss are designed to be the same in all four external branches. Specifically, each simulation theme is briefly described as follows: case (1) is designed for the pure loss process; case (2) is for the situation when the rate of gene proliferation is higher than that of gene loss; and case (3) is designed to evaluate the effect of short internal branch length, controlled by the constant *c*=1/2, 1/4, 1/8 and 1/16, respectively. In each case, the number of gene families at the root was set to be ranged from 50 to 1000.

**Table 2.** Simulation themes for genome phylogeny inference.

| | $\mu t$ | $\lambda t$ | $\alpha$ | $\beta$ |
| --- | --- | --- | --- | --- |
| **Case (1)** |  |  |  |  |
| **All branches** | 0.5 | 0 | 0 | 0.607 |
| **Case (2)** |  |  |  |  |
| **All branches** | 0.1 | 0.12 | 0.108 | 0.0901 |
| **Case (3)** |  |  |  |  |
| **External branches ($k=1, \dots, 4$)** | 0.1 | 0.12 | 0.108 | 0.0901 |
| **Internal branches ($k=5$ or $6$)** | $0.1c$ | $0.12c$ | $1.2b$ | $b$ |

Note: constant $c$ is scale to measure the shortness of internal branches; in our simulation study, we set $c=1/2, 1/4, 1/8$ and $1/16$ , respectively; the loss parameter of internal branches ( $b$ ) can be calculated by Eq.(2) for each $c$ value; and the proliferation parameter is always given by $1.2b$ .

In each simulation theme, 200 simulation replicates were generated. We used each methods (the infinite-state ML, the finite-state ML or Gu-Zhang-04 for the extended genome distance) to infer the phylogeny in each simulation replicate and then calculated the proportion of correct inference (Fig.3). Our main results are summarized as follows. First, as long as the internal branch length is not too short, shown by cases (1) and case (2), the correct percentages of all three methods were usually high as long as the number of gene families is more than 200. Second, in the case of short internal branch length, as exemplified by *c*=1/16 in case (3), the correct percentage was low but can be significantly improved when the number of gene families was increased to 500 or more. We thus concluded that the overall performance of finite-state ML was virtually the same as that of infinite-state ML.

**Fig. 3.**
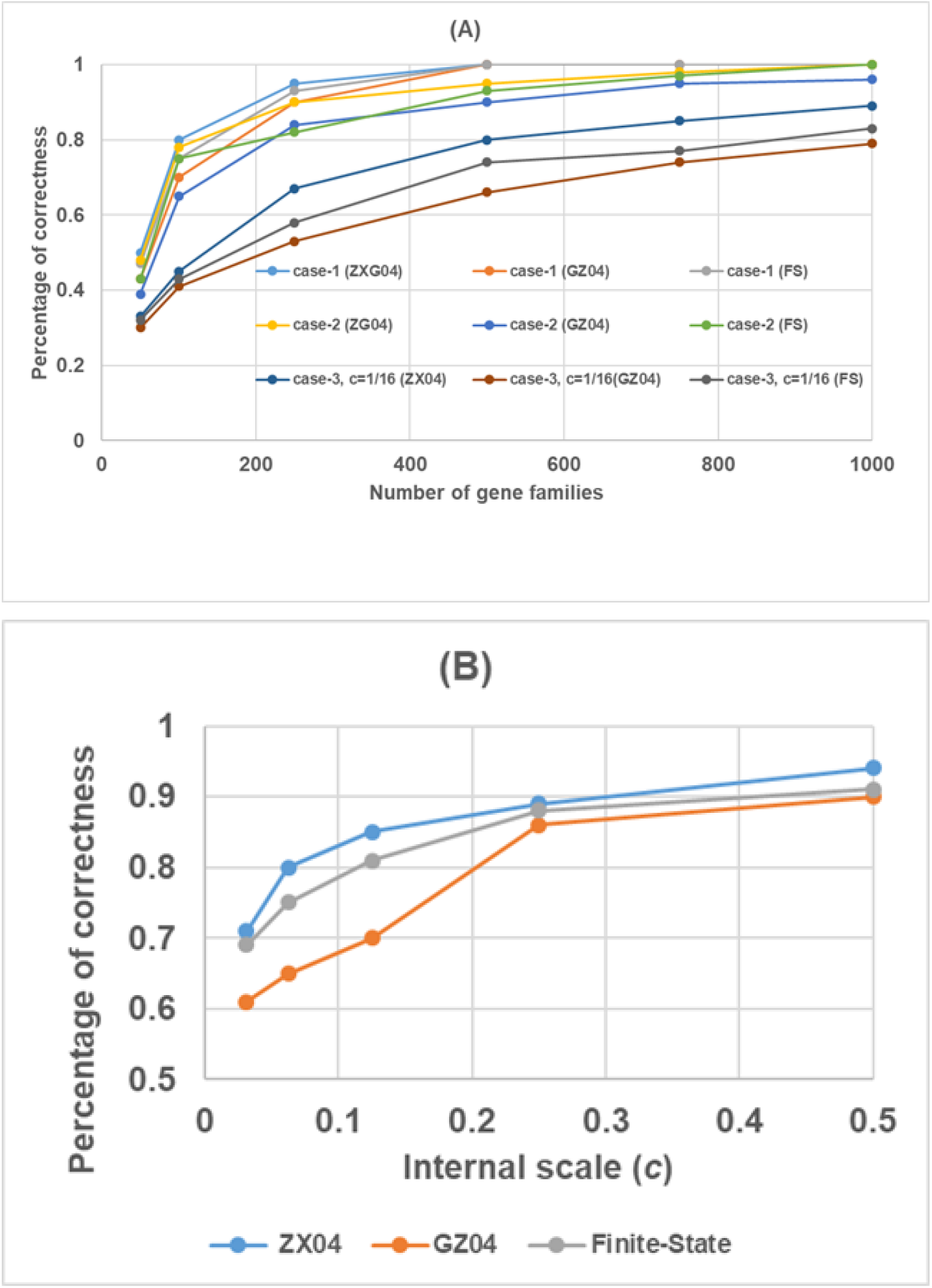
The percentage of correctness of genome phylogeny inference plotting against the number of gene families (panel A), or against the scale-constant of internal branch (panel B). The detail of simulation themes is described by Table 2. ZG04 short for the infinite-state ML method developed by Zhang and Gu (2004); GZ04 short for the distance method developed by Gu and Zhang (2004); and FS short for the finite-state ML developed by the current study.

### Case study: the universal genome tree of life

As a proof of concept, we applied the finite-state ML approach to infer a phylogenetic tree of four genomes from the COG database (http://www.ncbi.nlm.nih.gov/COG/): one Archaea, *Archaeoglobus fulgidus* (Afu), one Eukaryota, *Saccharomyces cerevisiae* (Sce), and two Bacteria, *Synechocystissp* and *Helicobacter pylori* (Syn and Hpy). In total 2785 gene families were used in this study. We maximized the likelihood function using the Newton-Raphson algorithm when a rooted phylogeny of the four species was given [11]. Among fifteen possible topologies, we first considered three different unrooted trees, each of which includes five related rooted trees (Fig.4A). Fig.4B shows the range of maximum log-likelihoods (logML) of rooted trees with the same unrooted tree, respectively. Those logMLs are overlapping between three unrooted trees, indicating that the root location of a phylogeny should be taken into account during the genome phylogeny inference.

**Fig. 4.**
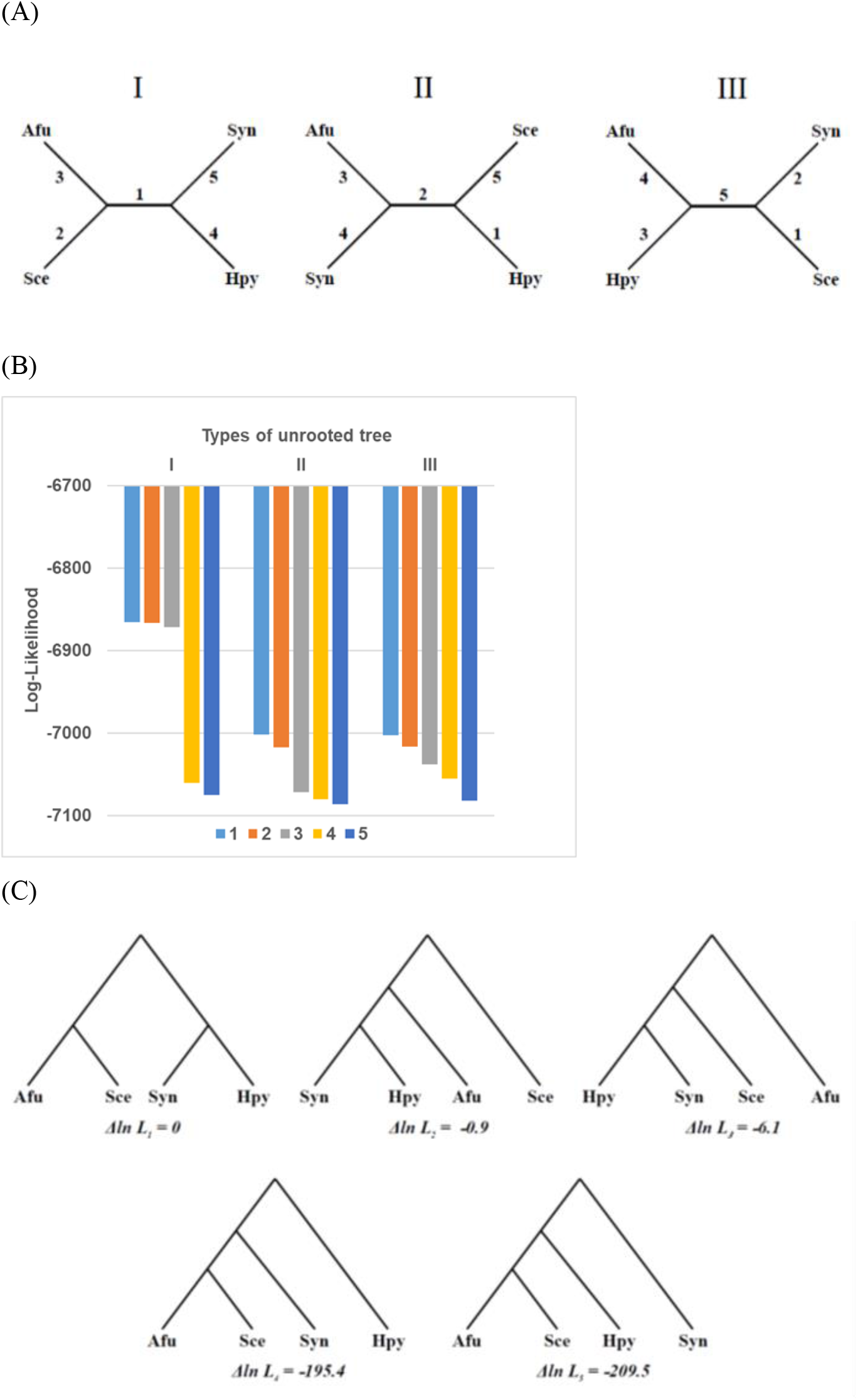
The finite-state ML phylogenetic analysis of four genomes: one Archaea, *Archaeoglobus fulgidus* (Afu), one Eukaryota, *Saccharomyces cerevisiae* (Sce), and two Bacteria, *Synechocystissp* and *Helicobacter pylori* (Syn and Hpy); 2785 gene families were used. (A) Three unrooted tree indicated by five possible rooting positions. (B) The range of the log-ML among fifteen possible topologies, represented according to three types of unrooted tree. (C) The logML differences for five rooted trees under the unrooted tree ((Afu, Sce), (Syn, Hpy)): ΔIn*L*_*k*_ = *logML*_*k*_ − *logML*_1_, *k*=1,…,5.

Moreover, we noticed that the largest logML value in ((Afu, Sce), (Syn, Hpy)) is considerably greater than those of other unrooted trees, respectively (Fig.4B). Fig.4C shows the logML differences for five rooted trees under the unrooted tree ((Afu, Sce), (Syn, Hpy)). It appears that the ML inferred genome tree supports the concept of universal tree of life: it groups the Bacteria (Syn and Hpy) together, and separates Bacteria from the Archaea (Afu) and the Eukaryota (Sce).

## Discussion

While vast amount of complete genomes are now available, a fast algorithm such as genome distance approach is useful to infer the genome phylogeny based on the J=1 or J=2 order gene contents [10, 11, 13, 16-20]. However, Gu [14] has shown that, though the genome distance can be accurately estimated under the pure birth-death process, the distance estimation could be unreliable when the new-gene input is considered. This finding indicated a limitation of the pairwise distance analysis, and a maximum likelihood approach is then highly recommended to explore the complexity of genome evolution. The problem is that, as shown by the work of Zhang and Gu [11], the maximum likelihood approach is computationally difficult so that it can only be applicable for a small number of genomes. The finite-state ML we developed here provides a plausible solution for the ML-based genome phylogeny analysis.

Future study will be focused on the feasibility of the finite-state ML for a fairly large number of genomes. To this end, we have to optimize the computation algorithm. Given the parameters, the computational time for calculating the likelihood function along a phylogeny is similar to that for the sequence-based likelihood. Moreover, a number of well-developed algorithms in molecular phylogeny to manipulate topologies during the ML search can be applied with some technical modifications. Our case-study indicated that the local ML problem could be the result of incorrect rooting. Since the possible number of rooted trees is much larger than unrooted tress, we need a new, more efficient strategy of topology search.

A difficult issue in practical implementation is how to efficiently deal with the complexity of parameter space, because the transition probabilities are much more complicated than the conventional Markov-chain model. Tentatively, I propose the following procedure that might be helpful. (*i*) Estimate the loss distance for each pair of genomes, because Gu [14] showed that the loss distance is insensitive to the existence of new-gene input. (*ii*) Infer the unrooted genome phylogeny based on the loss distance matrix by the neighbor-joining algorithm. (*iii*) Use the finite-state ML to determine the root under this unrooted phylogeny. (*iv*) Estimate the proliferation parameters and the new-gene input parameters under the rooted inferred phylogeny; one may construct different evolutionary scenarios, and then conduct a series of maximum likelihood ratio tests (see the discussion below). And (*v*) after carefully specified the parameter set, carry out the algorithm to search the maximum likelihood phylogeny.

Another challenging issue is to what extent one can utilize generalized gene contents to explore the pattern of genome evolution. The hierarchical LRT, or hLRT, is a classic statistical hypothesis testing of relative goodness of fit based on the statistic *Δ* = 2(ln*L*_1_ − ln*L*_0_), where *L*_0_ is the likelihood under the null hypothesis (simple model) and *L*_1_ is the likelihood under the alternative hypothesis (complex model). When the simpler model is a special case of the complex model, the significance of the *δ* statistic is usually assessed assuming it to be asymptotically distributed as *χ*2 with *q* degrees of freedom, where *q* is the difference in number of free parameters between the two models [23]. In cases where the null model has the parameter being tested fixed at a boundary of its parameter space, a mixed *χ*2 distribution was used to assess significance [24, 25]. One may anticipate that the implementation of hLRT is straightforward for a nested global hierarchy. For instance, *L*_*0*_ is the pure birth-death model, whereas *L*_*1*_ is the birth-death-input model.

To evaluate all the considered models (pure loss, pure birth, birth-death, and birth-death-input), one may implement this test successively performing pairwise comparisons of nested models in a hierarchical way [25]. [26] recognized there is no justification for preferring any one of the multiple options of sequence of parameter addition (or removal). It is possible that multiple models are selected as optimal by different sequences of parameter addition of the alternative hLRT. Alternatively, some other model-selection criteria have been proposed such as those based on the AIC [27] or Bayesian methods (BIC) [28-30]. Overall, those discussions suggested that caution should be taken into consideration when applying some standard hLRT protocols for model selection in the genome phylogeny analysis. In the case when the hLRT approach is used, it would be advisable to investigate as thoroughly as possible the impact of the hLRT scheme choice on the phylogenetic relationships of the species under study.

## Acknowledgements

The author is grateful to all members of the research group for constructive comments in the early version of this manuscript. Hongmei Zhang assisted the simulation study, and Zhan Zhou assisted the software development. The codes for simulation and case study are available upon the request.

## Conflict of interest

The author declares that he has no known competing financial interests or personal relationships that could have appeared to influence the work reported in this paper.

## Notes

### Competing Interest Statement

The authors have declared no competing interest.

